# *In silico* Neutron Relative Biological Effectiveness Estimations For Pre-DNA Repair And Post-DNA Repair Endpoints

**DOI:** 10.1101/2025.10.07.680965

**Authors:** Nicolas Desjardins, John Kildea

## Abstract

A comprehensive understanding of the energy-dependent stochastic risks associated with neutron exposure is crucial to develop robust radioprotection systems. However, the scarcity of experimental data presents significant challenges in this domain. Track-structure Monte Carlo simulations with DNA models have demonstrated their potential to further our fundamental understanding of neutron-induced stochastic risks. To date, most track-structure Monte Carlo studies on the relative biological effectiveness (RBE) of neutrons have focused on various types of DNA damage clusters defined using base pair distances. In this study, we extend these methodologies by incorporating the simulation of non-homologous end joining (NHEJ) DNA repair in order to evaluate the RBE of neutrons for misrepairs. To achieve this, we adapted our previously published Monte Carlo DNA damage simulation pipeline, which combines condensed-history and track-structure Monte Carlo methods, to support the standard DNA damage (SDD) data format. This adaptation enabled seamless integration of neutron-induced DNA damage results with the DNA Mechanistic Repair Simulator (DaMaRiS) toolkit. Additionally, we developed a clustering algorithm that reproduces pre-repair endpoints studied in prior works, as well as novel damage clusters based on Euclidean distances. The neutron RBE for misrepairs obtained in this study exhibits a qualitatively similar shape as the RBE obtained for previously reported pre-repair endpoints. However, it peaks higher, reaching a maximum RBE value of 23(1) at a neutron energy of 0.5 MeV. Furthermore, double-strand break (DSB) pairs with a separation distance of 11 nm better matched the RBE for misrepairs than did the DSB clusters defined with base pair distances that were used in previous *in silico* neutron RBE studies. These results suggest that there are some features to the spatial distribution of neutron-induced DNA damage that are best characterized either by explicit repair mechanism simulations or by cluster analysis based on Euclidean distances.

## 1 Introduction

### 1.1 Motivations

Neutron radiation exposure is a potential concern in radiotherapy with high-energy photon beams (Howell *et al* 2006), in particle therapy (Schneider and Hälg 2015), and in certain occupational environments such as in air travel (Goldhagen 2000) and in space travel (Stricklin *et al* 2021). The stochastic risks associated with neutron exposure are known to be highly energy dependent (ICRP 2003); however, quantification of this energy dependence is challenging due to the scarcity of suitable human exposure data sets (Ottolenghi *et al* 2013).

Practically, the stochastic risks of neutrons are quantified by scaling the better-known risks associated with x-rays and gamma rays, for which there is a more substantial body of literature. This approach assumes that the effects of neutrons and photons are qualitatively the same but quantitatively different. In the modern internationally-established system of protection against radiation exposure, as promulgated by the International Commission for Radiological Protection (ICRP) and the US Nuclear Regulatory Commission (NRC), these scaling factors are the radiation weighting factor *w*_R_ (ICRP 2007) and the quality factor *Q* (US NRC 2021), respectively. The values of *w*_R_ and *Q* for neutrons were agreed on by their respective agencies based on the results of various radiobiological studies using the relative biological effectiveness (RBE) formalism.

The RBE is an endpoint-specific metric that compares the effect of a given radiation quality, defined by its particle type and energy, to that of a reference radiation, which is typically 250 kVp x-rays, ^60^Co *γ*-rays, or ^137^Cs *γ*-rays . The RBE of a radiation quality *i* is defined as the dose ratio

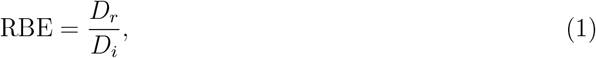

where *D*_*r*_ is the dose of a reference radiation, and *D*_*i*_ is the dose of a radiation quality of interest *i* that produces the same biological endpoint (e.g., number of chromosome aberrations) (CIRRPC 1995).

For simplicity, *w*_R_ and *Q* are assumed to be independent of endpoint, dose, and dose rate, and the factors for photons of all energies are set to unity. This; however, does not reflect the complexity of neutron RBE, which depends on those aforementioned quantities. For certain biological endpoints, the non-linearity of the photon dose–response curve also implies that the RBE of neutrons depends on both dose and dose rate. Furthermore, the biological efficacy of photons is weakly energy dependent, which complicates the comparison of RBE results based on different reference photons.

Ultimately, the difference in biological effectiveness of different radiation qualities for the same absorbed dose is understood to be the result of the difference in their energy deposition patterns. In the case of neutrons and photons, which are both indirectly ionizing particles, this translates into a difference in the energy deposition patterns of their respective charged secondary particles.

In human tissue or, even simpler, in a 4-element tissue equivalent material (Lund *et al* 2020), the secondary particle field of neutrons varies widely with neutron energy. The ratios of the particle types within the field change, the spectra of those particles change, and the relative dose contributions of each particle type change. On the scale of a human being, this also leads to significant variation of neutron RBE with depth as a result of continuous neutron moderation.

### 1.2 Prior work

Various studies (Baiocco *et al* 2016, Lund *et al* 2020, Montgomery *et al* 2021, Manalad *et al* 2023, Mentana *et al* 2025) have demonstrated that Monte Carlo simulations can reproduce key features of the energy dependence of neutron-induced stochastic risks from first principles. These studies employed a methodology that couples condensed-history and track-structure simulations (Baiocco *et al* 2016, Lund *et al* 2020, Montgomery *et al* 2021, Manalad *et al* 2023, Mentana *et al* 2025). Condensed-history Monte Carlo (CHMC) simulations make it computationally feasible to capture changes in the secondary particle field of neutrons at the scale of a human, and to do so in materials other than water, to which track-structure Monte Carlo (TSMC) simulations are otherwise typically confined (Lund *et al* 2020). In contrast, TSMC simulations provide the level of detail necessary to investigate energy deposition at subcellular scales.

More concretely, the methodology employed in these studies involved using CHMC simulations to determine the relative dose contributions and energy spectra of the various secondary particle species produced by neutrons, as a function of neutron energy and depth in the medium. The secondary particle spectra were then sampled to perform TSMC simulations aimed at investigating various endpoints for neutron RBE estimation. Lund *et al* (2020) provided RBE estimates based on the mean lineal energy, Montgomery *et al* (2021) and Manalad *et al* (2023) provided RBE estimates for explicit DNA lesions, and Baiocco *et al* (2016) did both.

The aforementioned three studies that scored DNA lesions used molecular DNA models with scoring units representing the individual components of nucleotides: the phosphate group, sugar and nitrogenous base. Together, these studies provided RBE estimates for various types of DNA lesions that are highly likely to result in misrepairs, which are considered an early step in the development of carcinogenesis (Jeggo and Löbrich 2007).

Another notable study is that of Zabihi *et al* (2020), which also investigated neutron-induced DNA damage with TSMC. Importantly, Zabihi *et al* (2020) investigated the interaction of two DSBs to simulate the formation of misrepairs. Their simulations; however, did not include indirect action of radiation; that is, the interaction with DNA of the free radicals produced by radiolysis, which was included in the work of Manalad *et al* (2023) and Baiocco *et al* (2016).

### 1.3 Objective 1

The first objective of this study was to compare how the neutron RBE for misrepairs resulting from both the direct and indirect action compares to that of the pre-repair endpoints presented in the works of Baiocco *et al* (2016) and Manalad *et al* (2023), which looked at various types of DSB clusters defined using base pair distances.

In order to achieve this goal, we adapted our previously published neutron-induced DNA damage simulation pipeline (Manalad *et al* 2023) to output the Standard DNA Damage (SDD) Data Format (Schuemann *et al* 2019b), making it compatible with the DNA Mechanistic Repair Simulator (DaMaRiS) toolkit (Warmenhoven *et al* 2019).

Furthermore, we wrote a Python script that reproduces the pre-repair endpoints presented in the works of Baiocco *et al* (2016) and Manalad *et al* (2023) from the SDD file in order to compare the RBE for misrepairs and pre-repair endpoints while keeping everything else consistent.

### 1.4 Objective 2

Two DSBs can be many base pairs away from each other or even on different chromosomes, but because of the high level of compaction of the DNA, they can still be within short Euclidean distances of each other, making the DSB pair a potential source of misrepair. Motivated by this, the second objective of this study was to determine the RBE for DSB clusters based on Euclidean distances. To achieve this, the aforementioned Python script also computes the yield of this new endpoint from the SDD file.

### 1.5 RBE for linear dose responses

We end this Introduction with a few words on the RBE formalism. Provided that both the radiation type of interest and the reference radiation have a linear dose response for the endpoint and dose range relevant to the RBE calculation, the RBE is a dose independent quantity. For linear dose response, we have isoeffect when

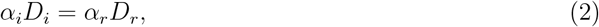

where *α*_*i*_ and *α*_*r*_ are the linear coefficients of the dose responses of the radiation type of interest and the reference radiation respectively. From (1) and (2) we get

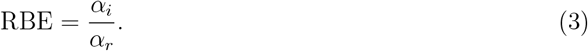

Conveniently, in the case where *D*_*i*_ = *D*_*r*_ = *D*, then (3) can be rewritten as

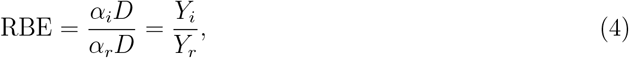

where *Y*_*i*_ and *Y*_*r*_ are the yields of the endpoint of interest. This quantity is typically referred as the radiation effect ratio (RER) and is equivalent to the RBE for linear dose responses. Given that the dose response of all endpoints of interest for this study were found to be linear within the dose range of interest, that is, at least up to 20 Gy, we obtained the RBE by computing the RER for 1 Gy, consistent with the work of Baiocco *et al* (2016), Montgomery *et al* (2021), Manalad *et al* (2023) and Mentana *et al* (2025). We did not carry simulations beyond 20 Gy but we can expected deviation from linearity at very high doses when the number of two track events becomes substantial.

## 2 Methods

### 2.1 Overview

The simulation pipeline can be broken down into four components. The first component is a series of CHMC simulations conducted to acquire the energy and relative dose contributions of the secondary particles released in tissue by neutrons and photons, see Figure 1 a). The second component is a series of TSMC simulations with a cell nucleus model performed to score various basic DNA lesions in the SDD format, see Figure 1 b). The resulting SDD files are then directly processed by a Python script to obtain pre-repair endpoints, see Figure 1 c), or fed into DaMaRiS to obtain misrepair outcomes, see Figure 1 d).

**Figure 1:**
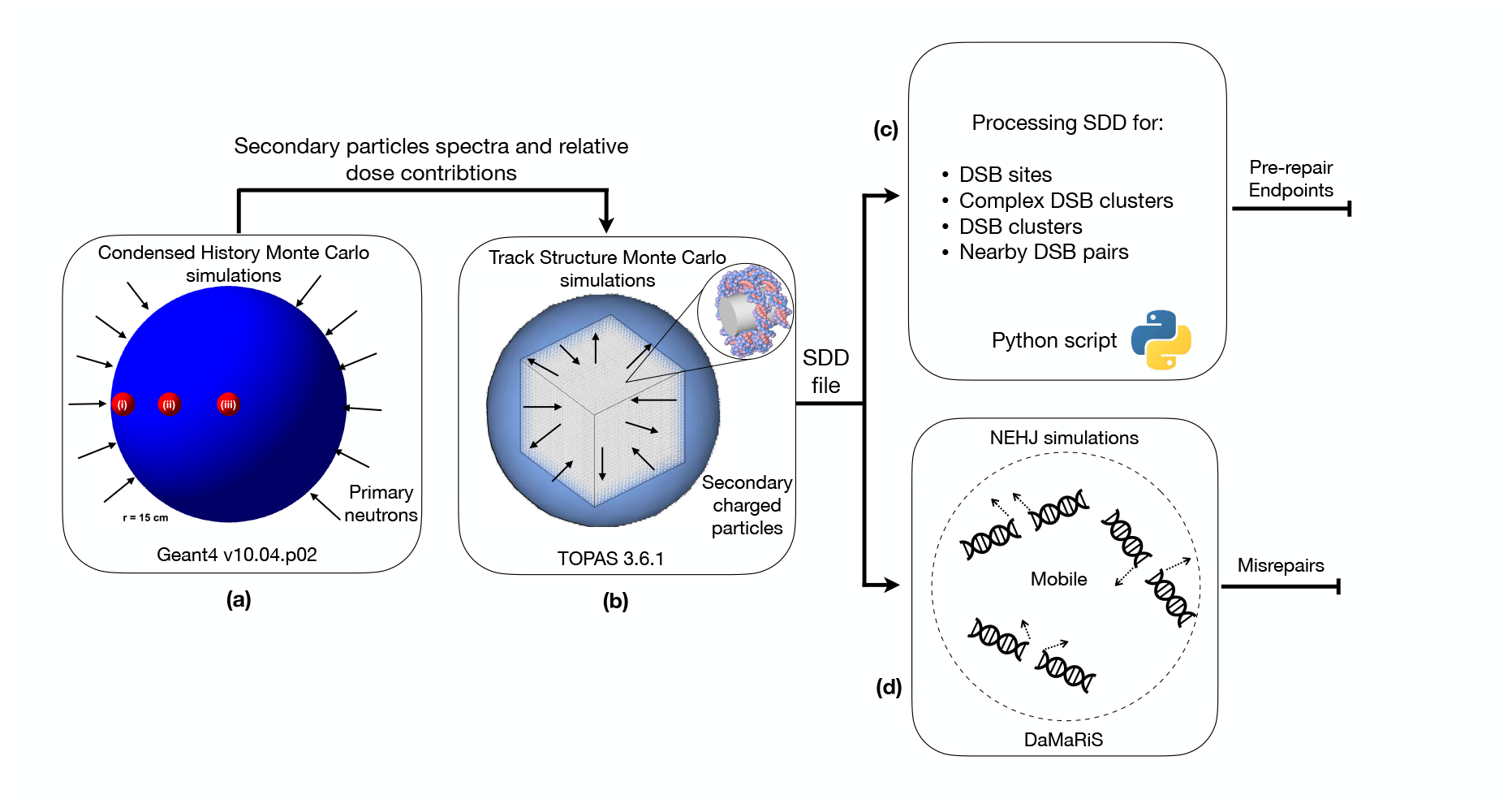
a) CHMC component with a 30 cm ICRU-4 sphere. The smaller red spheres are the scoring volumes. Only the outer scoring volume (i) is used for this work it has a radius of 1.5 cm and is centered 1.5 cm away from the surface. b) TSMC component with a nucleus model. The source is obtained sampling the secondary particle spectra from a). c) Python script that computes the pre-repair endpoints yields from the SDD file. d) Depiction of the mobile DSB ends in the spherical boundary with a diamater of 13.51 µm. The figure of the ICRU-4 sphere a) and the nucleus model with the octameter in the zoomed circle are modified illustrations reproduced from Montgomery *et al* (2021) ©2021 Institute of Physics and Engineering in Medicine. All rights reserved.

### 2.2 Condensed-history simulations

This aspect of the methodology has been detailed in our previous publication (Lund *et al* 2020), so we provide only a brief overview here. The goal of the CHMC simulations was to characterize the spectra and relative dose contributions of the various secondary particles generated by primary neutrons and photons as they pass through biological tissues. The simulations were carried with monoenergetic neutron sources, covering 18 energies ranging from 1 eV to 10 MeV, and a single photon source of 250 keV, which served as the reference energy for our RBE calculations. These simulations were conducted using Geant4 v10.04.p02 and its radiobiology extension Geant4-DNA (Incerti *et al* 2010a) with an ICRU-4 sphere (White *et al* 1989), which has a diameter of 30 cm.

Because the spectrum and the relative dose contribution of each secondary particle species is expected to change with depth in the medium, in the work of Lund *et al* (2020) three depths were sampled in the ICRU-4 sphere. However, only the outer scoring volume (with a diameter of 3 cm centered at a depth of 1.5 cm from the ICRU-4 sphere’s surface) will be considered in this study as it provides the maximal RBE. This aligns with the conservative approach adopted by the radiation protection agencies.

The secondary particle species released by neutrons are numerous but because of limitations in the current Geant4-DNA physics list for the required energy range (Incerti *et al* 2016), only the secondary electrons, protons, and alpha particles were considered in the next step. The relative dose contribution of those three particle species were renormalized to sum to unity.

### 2.3 Radiation-induced DNA damage simulations

The TSMC DNA damage simulations were conducted using the DNA model and a modified version of the scorer employed in our previous studies on complex DSB lesions (Montgomery *et al* 2021, Manalad *et al* 2023). The scorer, which was made as a TOPAS (Perl *et al* 2012) class, was modified to output DNA lesions in the SDD Format; however, all parameters pertaining to scoring lesions were left unmodified; therefore, the previous benchmarking conducted with mononenergetic protons and presented in Manalad *et al* (2023) remains applicable to this work.

Compared to our previous work (Montgomery *et al* 2021, Manalad *et al* 2023), a different TOPAS source parameter was used to specify the neutron secondary particle spectrum. The setting (s:So/Example/BeamEnergySpectrumType = ‘Continuous’) was applied instead of (s:So/Example/BeamEnergySpectrumType = ‘Discrete’). This change allows particles to be generated anywhere within the energy bin width of the particle energy spectrum as opposed to only the edge of the bin. The effect of this change on the pre-repair RBE estimation was evaluated. Other relevant simulation parameters can be found in Table 1.

**Table 1:** Track-structure DNA damage simulations parameters

| Parameter | Values |
| --- | --- |
| Physics Module | G4EmDNAPhysics_hybrid2and4<br>Combination of G4EmDNAPhysics_option2<br>and G4EmDNAPhysics_option4 |
| Strand break energy threshold | 17.5 eV |
| Base lesion energy threshold | 17.5 eV |
| Probability of HO<sup>•</sup> to cause damage | 40 % |
| DSB max length | 10 bp |
| Target dose | 1 Gy |
| Number of repeated simulations | 100 for neutron secondaries, 950 for photons |

### 2.4 DNA repair simulations

For the DNA repair simulations, we used DaMaRiS, a software developed by the PRECISE group at the University of Manchester that has been integrated to TOPAS-nBio (Henthorn *et al* 2018, Ingram *et al* 2019, Warmenhoven *et al* 2019). Briefly, DaMaRiS is a flexible tool that allows one to explore various user-defined repair pathways by defining various repair stages with their associated time constants. The repair pathway dictates the possible progression of the simulation’s “objects”, such as “DSB end objects” and “synaptic complex objects.” For this study, we used the NEHJ pathway provided in TOPAS-nBio and described in Warmenhoven *et al* (2019), with the default allowed repair time of 24 h. We selected the NEHJ only pathway as opposed to the combined NEHJ and HR pathway because our nucleus model best represents the state of a cell in the G0/G1 phase where HR is not available. DaMaRiS confines the moving DSB ends within a spherical reflective boundary. To accommodate our cubic cell nucleus model (Montgomery *et al* 2021), we used its circumscribed sphere as the simulation boundary, that is, a sphere with a radius of 6.755 µm.

### 2.5 Pre-repair and post-repair endpoints

The RBE is highly dependent on the endpoint being investigated. As stated in the introduction, one goal of this study was to compute various pre-repair endpoints reported in the literature, allowing for comparisons across endpoints while keeping other factors, such as the DNA model and scorer, constant. Additionally, this approach enables the evaluation of similarities in outcomes between our model and other models for the same endpoints.

A description of all endpoints investigated in this study can be found in Table 2. The two new endpoints presented here are the misrepair and the nearby DSB pair. Misrepairs are obtained with DaMaRiS, they include any DSB end that joined to the DSB end from a different DSB site. The two wrongly joined ends can originate from DSBs from the same chromosome or from different chromosomes. Nearby DSB pairs are pre-repair endpoints computed from the SDD file. They are defined as two DSBs within a specified Euclidean distance, measured between their centerpoints. Again, those two DSBs can be on the same chromosome or on different chromosomes. One DSB can be part of more than one nearby DSB pair.

**Table 2:** Descriptions of the endpoints used for RBE estimation in this work.

| Endpoint name | Description | Minimum number of basic lesions |
| --- | --- | --- |
| Pre-repair endpoints |  |  |
| DSB site | Location of one DSB | 2 |
| Complex DSB lesion<br>(Montgomery *et al* 2021)<br>(Manalad *et al* 2023) | Site with a least one DSB and at least one other lesion of any kind where each lesion is within 40 bp of its closest neighbour | 3 |
| DSB cluster<br>(Baiocco *et al* 2016) | Site with at least two DSBs within 25 bp | 4 |
| Nearby DSB pairs | Two DSBs where the centerpoints of the two DSBs are within a specified Euclidean distance | 4 |
| Post-repair endpoints |  |  |
| Misrepairs | Two DSBs end originating from distinct DSBs that joined together due to a DNA repair mechanism. | 4 |

\* In (Montgomery *et al* 2021, Manalad *et al* 2023) this type of lesion is referred to as a complex DSB cluster. To avoid confusion with lesions that require at least two DSBs, called DSB clusters in Baiocco *et al* (2016), we refer to it as a Complex DSB lesion in this work.

For each neutron energy and for the reference radiation, we conducted the DNA damage simulations pertaining to each secondary particle species independently. Hence, we combined species-specific damage yields via a weighted sum. The yield *Y* for a primary particle *P* of a given endpoint is obtained as follows:

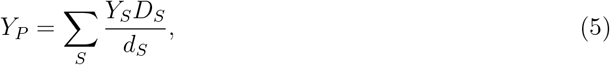

where *S* corresponds to the secondary particle species, *d*_*S*_ corresponds to the relative dose contribution of *S* obtained from the CHMC simulations, and *D*_*S*_ corresponds to the actual dose deposited by that species in a given TSMC simulation run. Note that for (*P* = neutron) we have (*S* = {*e*^−^, *p*^+^, *α*}) and for (*P* = photon) we have (*S* = {*e*^−^}). The neutron energy-dependent RBE(*E*) can be obtained using:

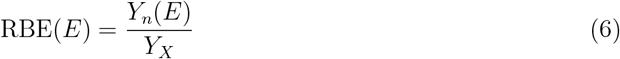

## 3 Results

### 3.1 Comparing pre-repair endpoints

Applying our Python clustering script to the SDD files resulting from our DNA damage simulations, we computed the RBE for the pre-repair endpoints investigated in Baiocco *et al* (2016), Manalad *et al* (2023), Mentana *et al* (2025), which are described in Table 2. In Figure 2, we compare our results to the RBE values from those studies. The endpoints a) DSB sites, b) Complex DSB lesions, and c) DSB clusters are presented in order of increasing minimum required basic lesions (see Table 2). The maximal RBE for each endpoint obtained in this study is a) 2.54(3), b) 4.78(8), and c) 16(1).

**Figure 2:**
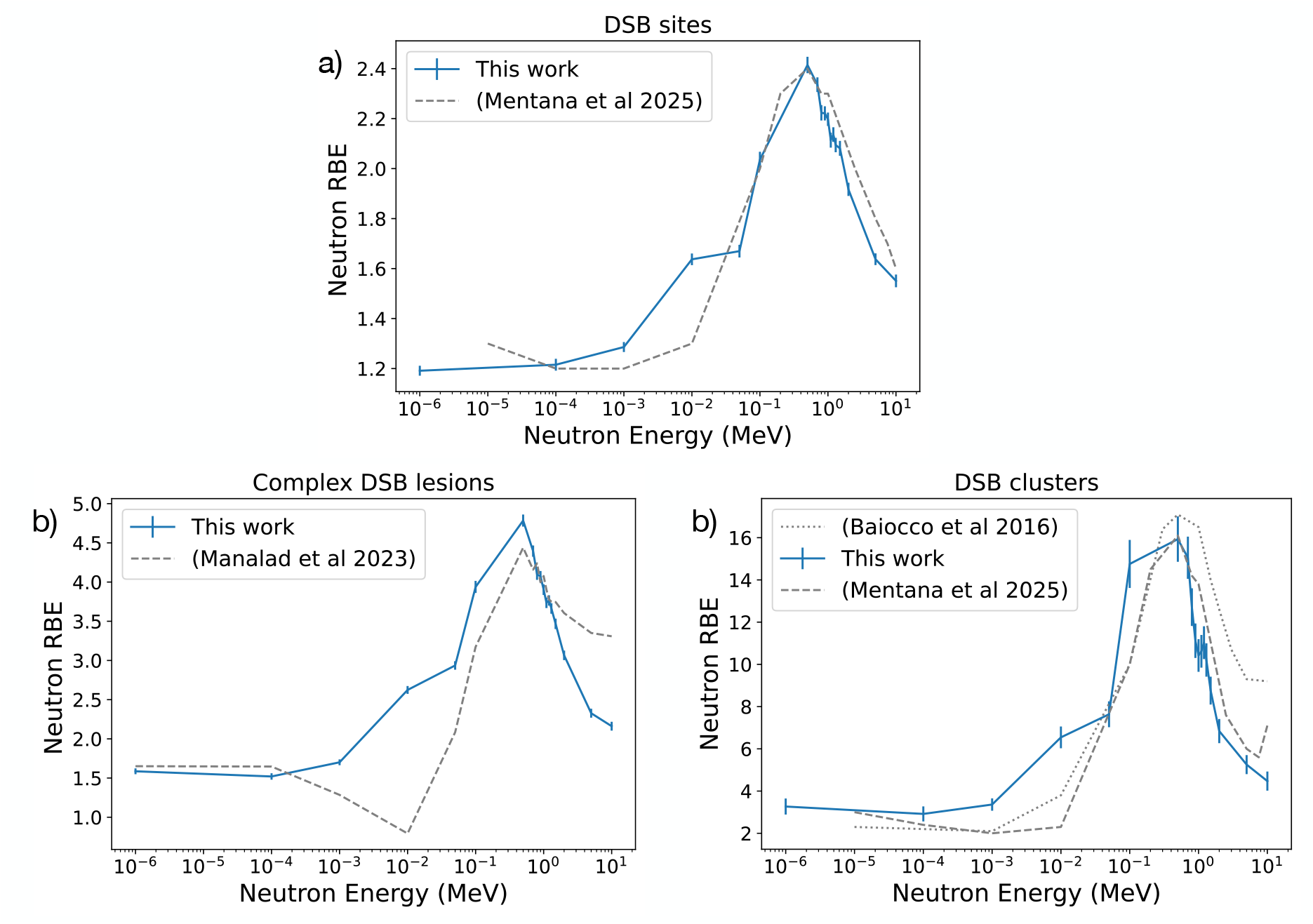
Comparison of neutron RBE values obtained in this study with literature results for pre-repair endpoints: (a) DSB sites, (b) Complex DSB lesions, and (c) DSB clusters. The three endpoints are detailed in Table 2. Results from (Baiocco *et al* 2016) were extracted using WebPlotDigitizer (Rohatgi 2024).

All error bars and quoted uncertainties were obtained through propagation of the standard uncertainty arising from the statistical variation of the simulations. In this study, for each neutron energy, 100 statistically independent simulations were used. The standard uncertainties were dominated by statistical variations across photon runs; therefore 950 photon simulations were conducted.

### 3.2 Misrepairs

Our SDD files were subsequently processed using DaMaRiS to compute the RBE for misrepairs. Only one misrepair RBE curve is presented in this study, as we did not vary the already benchmarked NEHJ parameters provided in TOPAS-nBio. In Figure 3 a), we compare the RBE for misrepairs to the RBE for the pre-repair endpoints obtained in this study. In Figure 3 b), we compare the RBE for misrepairs to *w*_R_, *Q*, and the RBE for DSB clusters from Baiocco *et al* (2016).

**Figure 3:**
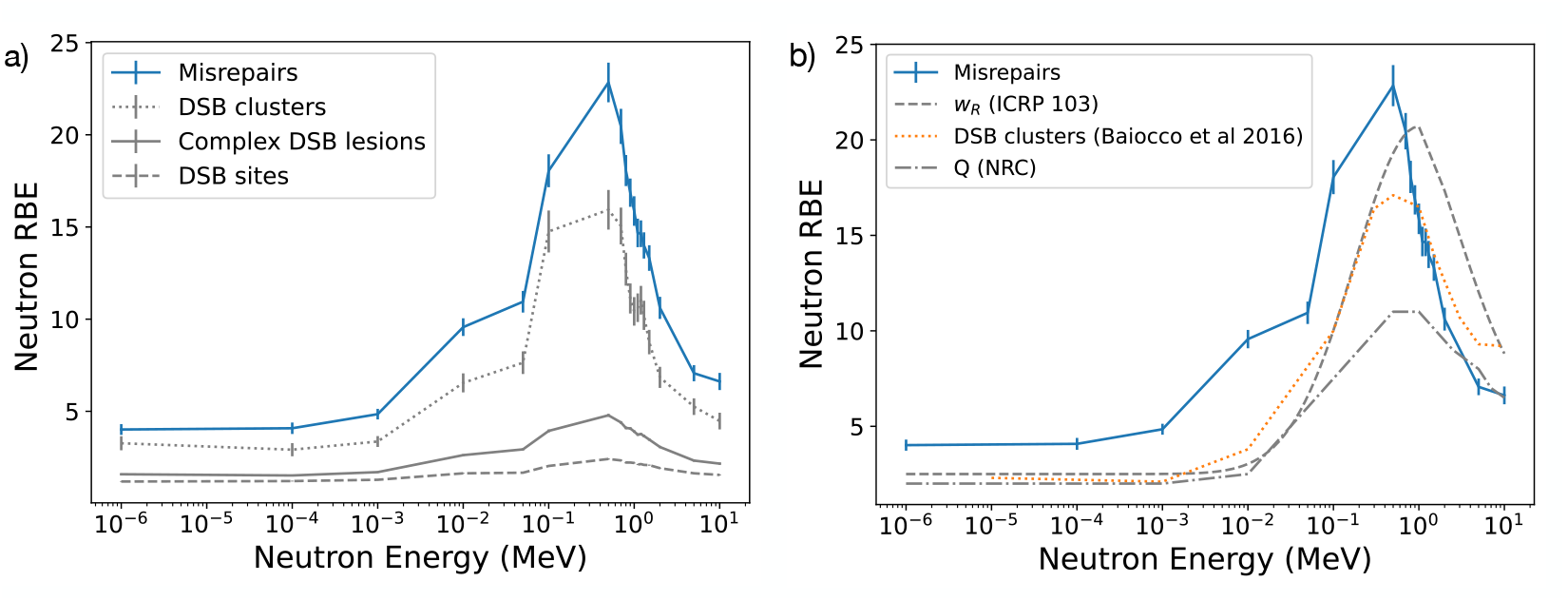
Misrepair results (a) Neutron RBE for misrepairs compared to the neutron RBE for various pre-repair endpoints. All the results in this panel were generated in this study. (b) Neutron RBE compared to the *w*_R_, *Q* and the RBE for DSB clusters from (Baiocco *et al* 2016). Values from (Baiocco *et al* 2016) were extracted using WebPlotDigitizer (Rohatgi 2024).

### 3.3 Nearby DSB pairs

The nearby DSB pair is a pre-repair endpoint, but the results pertaining to it are presented last because this endpoint was not used in prior work, and presenting the misrepair results first seemed more appropriate. The computed RBE for nearby DSB pairs varies as a function of the maximum Euclidean distance used to compute the number of DSB pairs. This dependence is shown in Figure 4 a) up to 80 nm for a neutron energy of 0.5 MeV, the energy at which the maximal RBE for nearby DSB pairs was observed.

**Figure 4:**
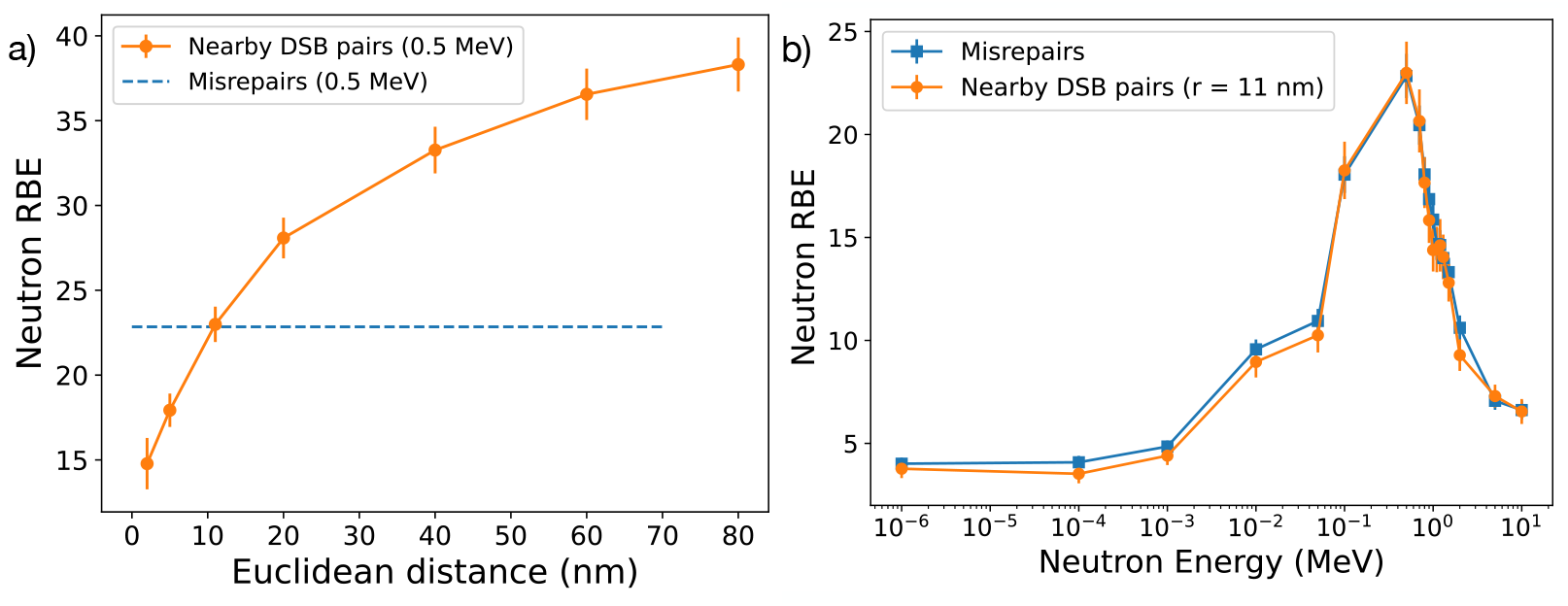
(a) Maximum RBE for 0.5 MeV neutrons as function of the Euclidean distance between nearby DSB pairs. b) RBE for misrepairs against the RBE for DSB pairs with a fix Euclidean distance of 11 nm.

While Figure 4 a) shows the variation of the RBE for nearby DSB pairs with Euclidean distance at a fixed energy of 0.5 MeV, Figure 4 b) shows the variation of the RBE for nearby DSB pairs with energy for a fixed Euclidean distance of 11 nm. The 11 nm Euclidean distance was selected to best match the misrepair values in the high RBE region of the curve. a) b)

## 4 Discussion

### 4.1 Comparing pre-repair endpoints across studies

In Fig. 2, we provide a comparison between the RBE curves obtained in this study and those obtained by Baiocco *et al* (2016), Manalad *et al* (2023), and Mentana *et al* (2025).

#### 4.1.1 *Comparison with* *Baiocco* et al *(2016)* and *Mentana* et al *(2025)*

In the work of Baiocco *et al* (2016) and their subsequent study (Mentana *et al* 2025), the CHMC simulations were carried out with the general purpose Monte Carlo particle transport code PHITS (Sato *et al* 2013) and the TSMC simulations with the biophysical simulation tool PARTRAC (Friedland *et al* 2011), using a different nucleus model than the one used in this study; hence, differences in the results are not unexpected. For the first step of the simulation pipeline, namely the CHMC simulations, the geometry used in this study was identical to that described in Baiocco *et al* (2016), consisting of spherical scoring volumes with a radius of r=1.5 cm. In the work of Mentana *et al* (2025), 1 cm-thick spherical shells were used instead, covering a narrower region by comparison. However, our results for both DSB sites and DSB clusters are in good agreement with those from Mentana *et al* (2025), and even better than with the results of Baiocco *et al* (2016) in the case of DSB clusters.

#### 4.1.2 *Comparison with Manalad* et al *(2023)*

The discrepancy between our previous RBE curve for complex DSB lesions presented in Manalad *et al* (2023) and the one presented in this study is entirely due to the *BeamEnergySpectrumType = “continuous”* parameter setting in TOPAS. Switching this parameter to *BeamEnergySpectrumType = “discrete”* yields the curve obtained by Manalad *et al* (2023), see Figure 5. For the TSMC simulations, the electrons, protons and alpha particles are sampled from energy distributions obtained with CHMC simulations. The parameter *BeamEnergySpectrumType = “discrete”* samples only the edges of the distribution bins, whereas the bins represent the likelihood of a range of energies and not a single point. This parameter change, which was justified *a priori*, makes the drop-off of the RBE curve steeper and results in better agreement with *w*_R_ and *Q*.

**Figure 5:**
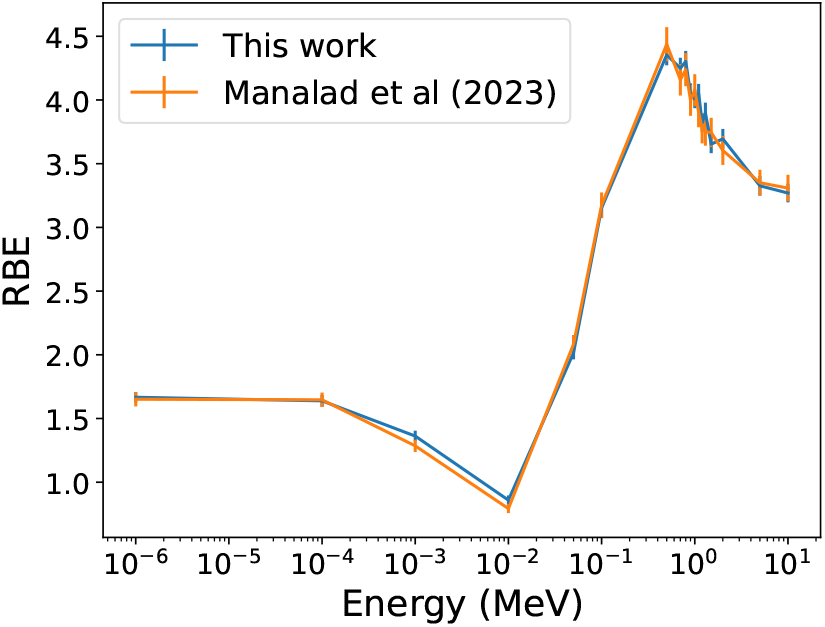
Comparison of the RBE curve for complex DSB lesions reported by Manalad *et al* (2023) with the curve obtained with the simulation pipeline used in this work, differing only by setting the TOPAS source parameter *BeamEnergySpectrumType* to “discrete,” as in Manalad *et al* (2023).

### 4.2 Comparing pre-repair endpoints obtained in this study

We now aim to compare the RBE associated with the various endpoints while maintaining all other parameters consistent. The definitions given to those endpoints allow them to have various numbers of basic lesions, but each must meet a minimum number of lesions which were listed in Table 2. For the endpoints investigated in this study the maximal RBE increases with the required minimum number of basic lesions suggesting this could be a determining factor. This pattern can be observed across panels in Figure 2.

One possible explanation for that correlation is that the most basic form of an endpoint is the most likely form. In that case, the overall yield would be dominated by that form and because neutrons yield more complex lesions than photons, the RBE would increase with increasing level of required minimal complexity. In other words, the idea suggested here is that even if the definition of an endpoint allows for very complex lesions, such as is the case with the Complex DSB lesion from Manalad *et al* (2023), it is its most basic form that will dictate the RBE and not how complex the endpoint can be.

### 4.3 Misrepairs

Peaking at an RBE value of 23(1) for neutrons of 0.5 MeV, the RBE for misrepairs exceeds the RBE of the complex DSB lesions, DSB clusters, and DSB sites. Misrepairs in this work include any joining of incongruent DSB ends. DaMaRiS cannot simulate the alterations in the base pair sequence that often result from NEHJ even when the correct DSB ends are joined. We decided not to distinguish among the different kinds of chromosomes aberrations either, because without a biologically accurate chromosome mapping the ratio between the different yields of chromosomes aberrations would be meaningless.

For the development of the DaMaRiS toolkit, DNA damage simulations were needed; therefore, the kinetics of the repair models used by DaMaRiS are inherently influenced by the DNA damage simulation used in its development. The effects of compensatory mechanisms in repair models due to the choice of DNA damage simulation tools are discussed at length in Warmenhoven *et al* (2023). Notably, the authors demonstrate that DNA damage simulation tools resulting in more clustered damage lead to repair models that restrict the range over which DSB ends interact. Two repair simulation tools can produce the same final results using different kinetics, provided they are coupled with the DNA damage tool with which they were benchmarked. To demonstrate how our DNA damage simulation tool compares to the one used to benchmark DaMaRiS, Figure 6 compares the benchmarking results obtained by Manalad *et al* (2023) to those obtained with the DNA damage simulation tool used in the development of DaMaRiS, as published in Warmenhoven *et al* (2023). The agreement between these results suggests that coupling our DNA damage model with the DaMaRiS toolkit should yield accurate outcomes.

**Figure 6:**
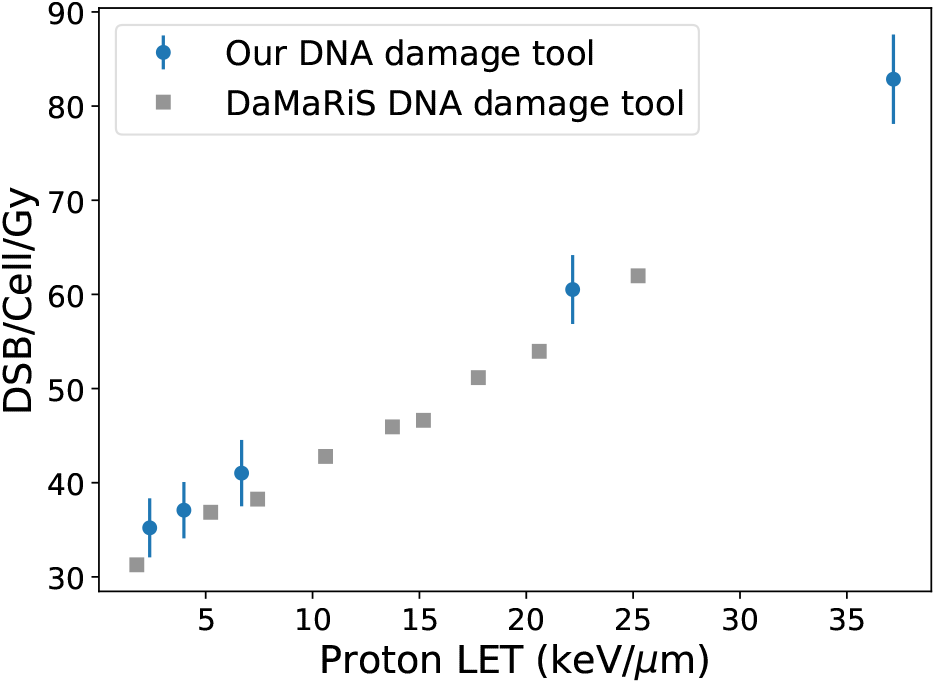
Comparing the DSB yield per cell per gray against proton LET for our DNA damage simulation tool (Manalad *et al* 2023) to the the the one used to benchmark DaMaRiS (Warmenhoven *et al* 2023). The data from (Warmenhoven *et al* 2023) were extracted using WebPlotDigitizer (Rohatgi 2024) and, therefore, are presented without error bars in this work.

### 4.4 Nearby DSB pairs

By varying the maximum Euclidean distance used to compute the nearby DSB pairs we found that 11 nm results in an RBE for nearby DSB pairs that best matches the RBE obtained for misrepairs. The strong agreement between the two curves, see Figure 4 b), suggests that nearby DSB pairs can be used as a good estimator of DSB misrepairs.

We will now seek to provide a rationale for the shape of the curve for the maximal RBE as function of Euclidean distances provided in Figure 4 a). To do so we introduce Figure 7, which is the same figure as Figure 4 a), except over Euclidean distances that span the whole nucleus. The y-axis is labeled RER in this case as opposed to RBE because over large distances the linearity of the dose response for nearby DSB pairs does not hold anymore, and therefore, the RBE and RER are not equivalent.

**Figure 7:**
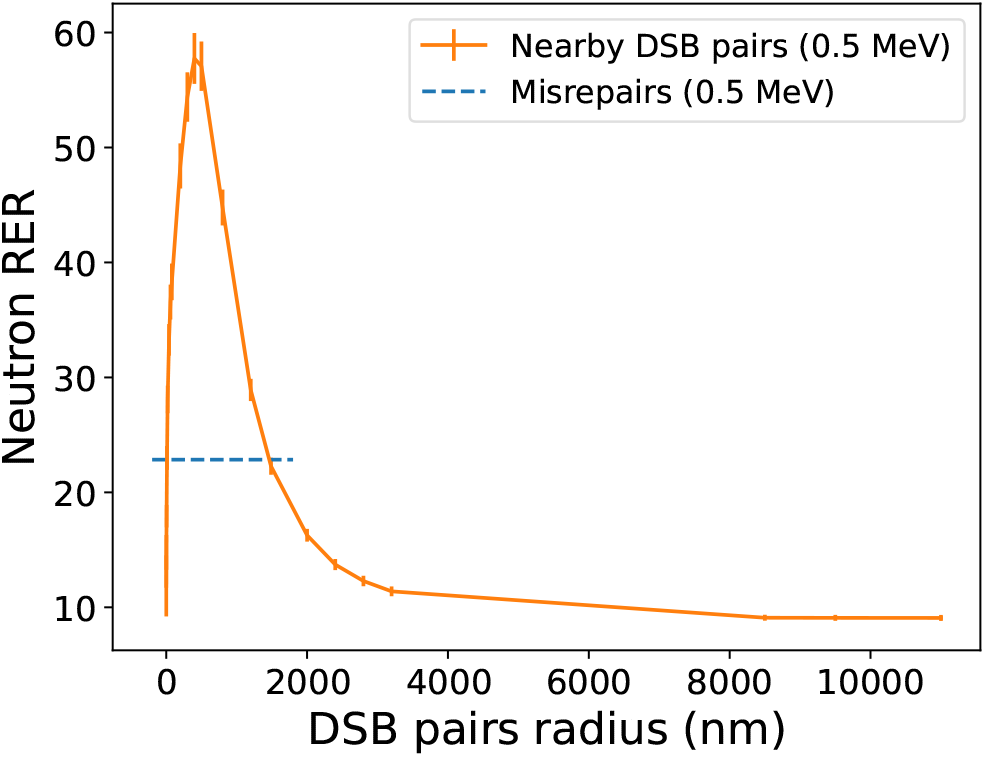
Neutron RER for nearby DSB pairs as function of the Euclidean distance for 0.5 MeV neutrons. This graph is shows how the RER behaves as a whole, but the values of the RER are not biologically relevant for this whole range of distances.

For a given irradiation condition, the variation in both the neutron and photon nearby DSB pair yields (*Y*_*n*_ and *Y*_*X*_) with respect to Euclidean distance follows sigmoid curves. The plateau of these sigmoid curves is given by the combination *C*(*n*_DSB_, 2), which represents the number of all possible pairs that can be formed from the DSBs present in the cell. The shape of Figure 7 is, therefore, the ratio of two sigmoids. For a single irradiation, the vertical asymptote seen in Figure 7 is given by the ratio of *C*(*n*_DSB_, 2)_*n*_ to *C*(*n*_DSB_, 2)_*x*_; however, this asymptote cannot be computed using the mean DSB yield from neutron and proton irradiation repeats, because *C*(*n*_DSB_, 2) is not a linear operation.

Therefore, the log-like appearance of Figure 4 a) corresponds to the first portion of the ratio of the two sigmoids. It is the stage where the yield for neutrons increases faster with respect to Euclidean distances than does the yield for photons. This also means that there is a second crossing point with the RBE for misrepairs; however, this crossing point, which occurs for a Euclidean distance around 1488 nm, is of no biological relevance.

### 4.5 Other remarks

The values of *w*_R_ and *Q* at very low neutron energies are approximately 2.5 for *w*_R_ and 2 for *Q*. In contrast, the RBE obtained for misrepairs at very low energies is around 4. This discrepancy could be partly explained by the fact that we are examining only the outer portion of the ICRU-4 sphere rather than RBE values for its entirety. In the work of Baiocco *et al* (2016), Montgomery *et al* (2021), and Manalad *et al* (2023), the outer scoring volume consistently shows the highest RBE at low neutron energies among all scoring volumes. Furthermore, some experimental studies on neutron RBE at low energies suggest that the current radioprotection system may underestimate neutron RBE in this range (Paterson *et al* 2022). Finally, the curves obtained in this work are endpoint-specific RBE curves and are therefore not directly comparable to *w*_R_ and *Q*, which were compiled from multiple types of data.

A major limitation for the study of neutron RBE with Geant4-DNA at the moment is the absence of models for low-energy heavy ion transport. Oxygen, nitrogen and carbon makes up 3.3 % of the dose at 0.5 MeV and 9.2 % of the does at 10 MeV (Lund *et al* 2020). However, this issue can be easily addressed once GEANT4-DNA physics lists capable of handling low-energy heavy ions are made available.

## 5 Conclusion

In this study, we employed Monte Carlo simulations to investigate neutron RBE for both pre-DNA repair and post-DNArepair endpoints. To the best of our knowledge, this represents the first *in silico* study to present RBE estimates for misrepairs, and DSB clusters based on Euclidean distances from both the direct and indirect effects of ionizing radiation. This was achieved by modifying our previously-published DNA damage simulation pipeline to ensure compatibility with the SDD format, enabling seamless integration with the repair simulation toolkit DaMaRiS with as well as with a novel in-house clustering algorithm.

The neutron RBE for misrepairs that we obtained exhibits a higher peak compared to previously published pre-repair endpoints. However, it is closely matched by the RBE for inducing nearby DSB pairs with a maximum distance of 11 nm. This suggests that neutron-induced DNA lesions may have an inherent structural organization that is better captured by explicit repair mechanisms or DNA clusters defined by Euclidean distances rather than by base pair distances along the genome.

This study lays the groundwork for refining simulation techniques, which will, in turn, enhance our fundamental understanding of the stochastic risks associated with exposure to neutron radiation. However, it does not yet provide a fully accurate assessment of neutron RBE, highlighting the need for ongoing research and development in this area.

## 6 Acknowledgements

We are grateful to Chris Lund, Logan Montgomery, James Manalad, and Felix Mathew for their invaluable contributions to this research project. We extend special thanks to Chris Lund for providing access to the neutron secondary particle spectra, which is an integral part of this project’s pipeline. We also thank Logan Montgomery and James Manalad for sharing their TOPAS scorer and geometry, which were integral to the development of this work. Finally, we are thankful to Felix Mathew and James Manalad for their ongoing support and insightful feedback throughout the project, and to Luc Galarneau for laboratory administration support. This research was made possible in part through computational resources provided by Calcul Québec and the Digital Research Alliance of Canada.

Funding for this research was provided by the Collaborative Research and Training Experience grant entitled Responsible Health and Healthcare Data Science (SDRDS) of the Natural Sciences and Engineering Research Council, The Canada Foundation for Innovation John R. Evans Leaders Fund, the Canadian Space Agency Grant #19FAMCGB25 held by JK, a Discovery Grant of Natural Sciences and Engineering Research Council of Canada held by JK, and a Fonds de recherche du Québec – Nature et technologies Masters Training Award received by ND.

Additionally we gratefully acknowledge the following open-source softwares: Geant4 (Agostinelli *et al* 2003), Geant4-DNA (Incerti *et al* 2010b,c, Bernal *et al* 2015, Incerti *et al* 2018, Tran *et al* 2024), TOPAS (Perl *et al* 2012), TOPAS-nBio (Schuemann *et al* 2019a), DaMaRiS (Warmenhoven *et al* 2019), WebPlot Digitizer (Rohatgi 2024), NumPy (Harris *et al* 2020) and Matplotlib (Hunter 2007)

